# Sensitivity based model agnostic scalable explanations of deep learning

**DOI:** 10.1101/2025.02.21.639516

**Authors:** Manu Aggarwal, NG Cogan, Vipul Periwal

**Affiliations:** National Institutes of Health, Bethesda, MD; Department of Mathematics, Florida State University, Tallahassee, FL

**Keywords:** explainable AI, global sensitivity analysis

## Abstract

Deep neural networks (DNNs) are powerful tools for data-driven predictive machine learning, but their complex architecture obscures mechanistic relations that they have learned from data. This information is critical to the scientific method of hypotheses development, experiment design, and model validation, especially when DNNs are used for biological and clinical predictions that affect human health. We design SensX, a model agnostic explainable AI (XAI) framework that outperformed current state-of-the-art XAI in accuracy (up to 52% higher) and computation time (up to 158 times faster), with higher consistency in all cases. It also determines an optimal subset of important input features, reducing dimensionality of further analyses. SensX scaled to explain vision transformer (ViT) models with more than 150, 000 features, which is computationally infeasible for current state-of-the-art XAI. SensX validated that ViT models learned justifiable features as important for different facial attributes of different human faces. SensX revealed biases inherent to the ViT architecture, an observation possible only when importance of each feature is explained. We trained DNNs to annotate biological cell types using single-cell RNA-seq data and SensX determined the sets of genes that the DNNs learned to be important to different cell types.

## 1 Introduction

Deep neural network models (DNNs) are powerful tools for learning patterns in large data sets to make predictions with high accuracy. Further, formulation of DNNs does not require any explicit system-specific rules or constraints to be imposed a priori and can be purely data driven. This makes DNNs indispensable when there is a lack of knowledge of the system-specific mechanistic and functional relations. The success of DNNs is theoretically assured by their architecture of interconnected networks with a sufficiently large number of parameters [Hornik et al., 1989], which often ranges from millions to trillions in practice. The complexity of the interconnectedness of such a large number of parameters often obscures explanation of the mechanistic relations that a trained DNN has learned as the criteria for its predictions. However, this is crucial when they are used to make biological, and especially clinical, predictions. For example, DeGrave et al. [2021] showed that DNNs trained to detect COVID-19 in chest radiographs relied on spurious factors rather than medical pathology. Consequently, these models failed in new hospitals even though they had high accuracy on test data. Besides conceptual validation of what a DNN has learned, another important reason for explaining DNNs is that determining mechanistic relations is a critical step in the scientific method of developing hypotheses and designing experiments to further our scientific knowledge. This has the potential to connect successful data driven ‘black-box’ methods with traditional mechanistic models, substantially expanding our understanding of both approaches.

The field of research to understand the mechanisms underlying DNNs has been referred to as explainable AI (XAI) [Gunning and Aha, 2019, Arrieta et al., 2020, Linardatos et al., 2020] and also as model interpretability [Lipton, 2018]. However, there is no consensus on the definitions of explainability and interpretability [Lipton, 2018, Arrieta et al., 2020]. We adopt the broad definition of explainability proposed by Arrieta et al. [2020]—explainability is to produce post-hoc details or reasons of a trained model to make its functions clear and easy to understand. In this regard, we first illustrate what our framework explains about a model.

A DNN trained to identify smiling human faces predicts with a 99% confidence that the face shown in Figure 1A is smiling. The DNN is the deterministic mathematical model, the model input is the image of size 224 pixels by 224 pixels by 3 color channels, and the corresponding model output is the predicted confidence that the face in the image is smiling. We say that the number of features of the input or the number of dimensions of the input space is 224 *×* 224 *×* 3 = 150, 528. A random perturbation of all features results in a new image (Figure 1B) for which the model predicts smiling with a reduced confidence of 13%. *Perturbations in which features of the input were important to change the model’s prediction?* Our framework ranks the importance of features of a given input based on the sensitivity of the model output to perturbations in the features of the input.

**Figure 1:**
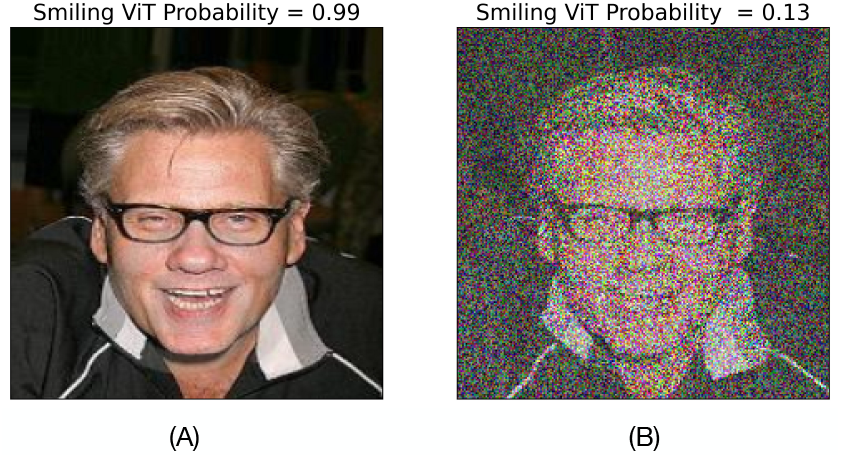
XAI based on global sensitivity analysis: (A) Vision transformer (ViT) trained to identify smiling human faces predicts that the face in the image is ‘Smiling’ with 99% confidence. (B) Perturbing features (all pixels across three color channels) reduces the model’s predicted confidence of ‘Smiling’ to 13%. *To perturbations of which input features is this change in the model confidence sensitive to?*

**Figure 2:**
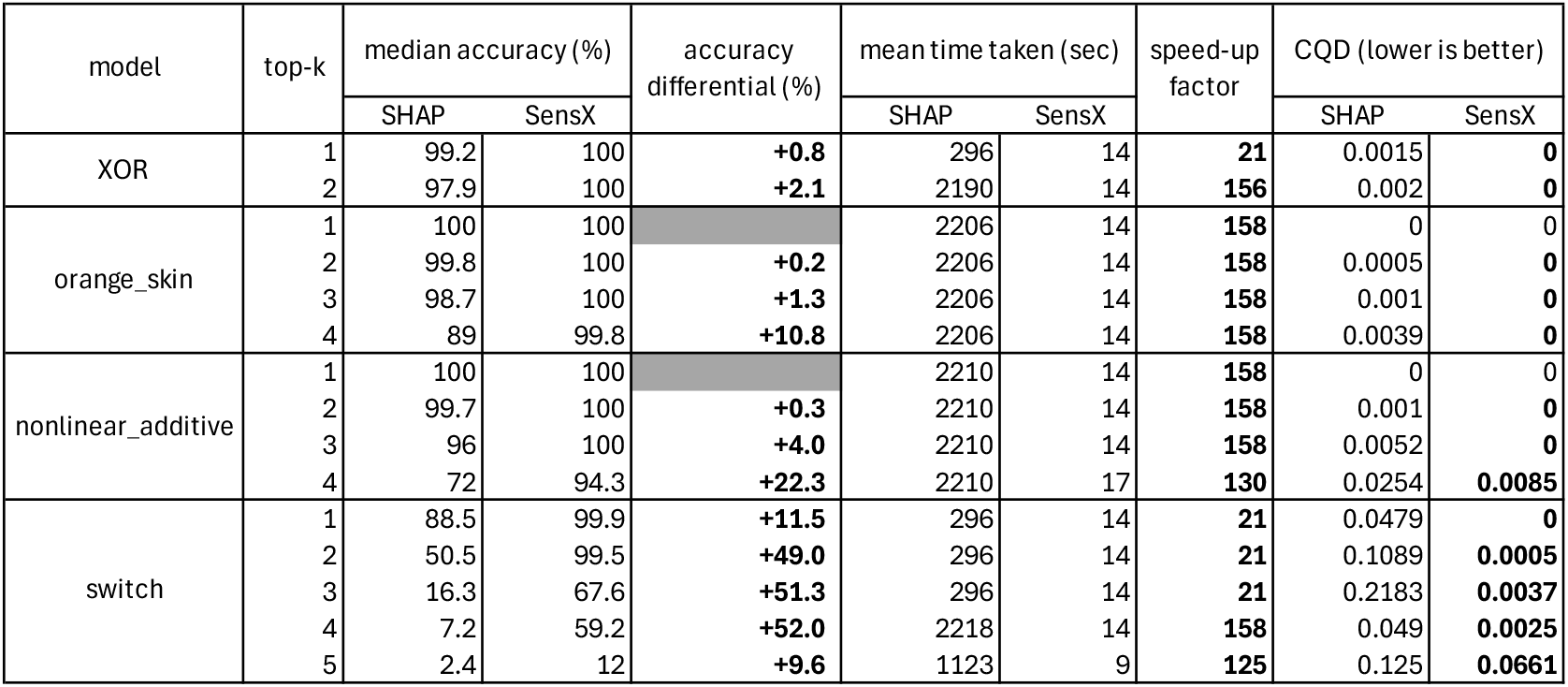
Benchmarks using synthetic data sets to compare SHAP and SensX for accuracy, computation time, and consistency. Consistency is compared using coefficient of quartile deviation (CQD), rounded to four decimal places.

This approach to XAI comes under the umbrella of sensitivity analysis (SA). In general, SA determines the effects of perturbations in input features on a scalar quantity of interest (QOI) that is a mathematical function of the features. SA methods are classified as local and global SA. Local SA considers perturbations in one input feature at a time keeping the rest fixed. Global SA (GSA) is a more comprehensive analysis because it considers simultaneous perturbations in all input features. However, GSA incurs a high computational cost. For example, Sobol’ indices [Sobol, 2001] are popular GSA measures [Tosin et al., 2020], but their computational cost limits their application to a few tens of features [Ikonen, 2016]. Van Stein et al. [2022] compare computational feasibility of different GSA methods for XAI (GSA-XAI), but the number of features was limited to less than 100. In comparison, the example shown in Figure 1 has hundreds of thousands of features. Nevertheless, Paleari et al. [2021] and Van Stein et al. [2022] empirically showed that the Morris method [Morris, 1991] is the least computationally costly GSA method among those tested, without compromising reliability and accuracy.

Besides the concern of computational scalability, a conceptual concern that we claim to be fundamental to GSA-XAI is to justify the domain of perturbations. To explain why, we introduce three criteria in this work to define justifiable perturbations as follows. First, we say that the perturbations should be contextually meaningful. For example, if the input features are gene expression values of different genes, then GSA based on perturbations that result in biologically infeasible gene expression values (negative, arbitrarily large, etc.) might not be reasonable. This is particularly important for XAI since many black-box DNNs are inherently mathematically defined for all real numbers and will generate an output given any input. Second, the perturbations should have a significant effect on the model output. Otherwise, the resulting GSA measures are not relevant and can be misleading. We denote these by significant perturbations. Third, since we attribute importance to features for a given model input, the perturbations should be with respect to that input. We denote these by grounded perturbations. We next introduce our framework and clarify grounded perturbations.

We design SensX, a model agnostic XAI framework that conducts GSA using justifiable perturbations to explain model inference for a given input. Our GSA is based on the Morris method which we briefly outline follows. A set of simultaneous perturbations in all features can be visualized as a trajectory in the input space, perturbing exactly one of the features along each step of the trajectory. Gradients are computed at each step using finite differences. When repeated for multiple sets of simultaneous perturbations, we get distributions of gradients with respect to features. Morris defined the mean and standard deviation of these distributions as linear and non-linear/interaction elementary effects of features, respectively. We implement and modify the Morris method as follows. First, the framework computes an input-specific domain of justifiable perturbations. We show later in one of our case studies that the input-specific domain might be much smaller than the global domain of the input space. This can potentially increase the accuracy of the results for the same number of model simulations since the elementary effects are estimated by sampling the domain. Second, the sets of simultaneous perturbations are generated such that the corresponding trajectories always start at the given model input. We say that they are *grounded* at a fixed reference point. In this work, we compute the mean of the distributions of absolute values of the gradients as suggested by [Campolongo et al., 2007] to get a single measure of importance for each feature. However, linear and nonlinear effects as proposed in the traditional Morris method can be computed without significant computational overhead because no additional model simulations are required.

Out of the vast repository of XAI methods [Linardatos et al., 2020], we compared SensX with SHapley Additive exPlanations (SHAP) [Lundberg, 2017] because it is the most widely used model agnostic XAI method (31, 309 citations according to Google Scholar, February 2025). Furthermore, it theoretically unifies multiple other methods, including LIME [Ribeiro et al., 2016] and DeepLIFT [Shrikumar et al., 2017]. Two other model agnostic XAI methods that we considered but did not pursue are L2X [Chen et al., 2018] and TERP [Mehdi and Tiwary, 2024]. We considered L2X because it was shown to outperform SHAP in accuracy and computation time for synthetic data sets. However, a required hyperparameter *k* of the method is defined as the number of important features, and its known true value was chosen to benchmark synthetic data sets in Chen et al. [2018]. It was not clear to us how the choice of *k* will affect the accuracy and computation time when its true value is not known, especially when there are hundreds of thousands of features. Arguably, L2X could be carried out for a set of chosen values of *k* when its true value is unknown, but we do not know how to decide on an optimal value of *k* based on the results of the different choices. We considered TERP because of its recency and unique explanations based on thermodynamics. However, preliminary simulations showed a higher computation cost and lower accuracy than SHAP for synthetic data sets.

We show three case studies. (1) We benchmark SensX and SHAP using four synthetic data sets. (2) We explain vision transformer (ViT) models that are trained to identify specific facial attributes in images of human faces. (3) We train feedforward neural networks (FNNs) for automated cell annotation using single-cell RNA-seq data sets and determine the sets of genes they have determined to be important to identify different cell types.

## 2 Results

We show results from three different case studies using different model architectures. (1) FNNs for synthetic data sets labeled 0 or 1, (2) vision transformer (ViT) models for images of human faces annotated for smiling and eyeglasses, and (3) FNNs for single-cell RNA-seq data annotated with 21 different cell types. All models are binary classifiers that are trained on labeled data. For multi-class data sets in (2) and (3), binary classifiers are trained separately to identify each class. The QOI for GSA of each model is the confidence predicted by the model that the input belongs to the class (predicted probability of labeling the input 1). We will refer to this as the model confidence.

### 2.1 Benchmarking SensX and SHAP

We benchmark SensX and SHAP for four synthetic data sets—XOR, orange skin, nonlinear additive model, and switch [Chen et al., 2018, 2017]. Binary-labeled ten-dimensional samples (or model inputs) were generated for each data set according to the criteria detailed in Methods. The exact set of relevant features is known for each data set from the labeling criteria. The number of relevant features is 2, 4, 4, and 5 for XOR, orange skin, nonlinear additive model, and switch, respectively. To highlight the input-specific analysis of both methods, the set of five relevant features can be different for different samples in the switch data set.

We computed the SensX and SHAP values for 1000 test samples and ranked the features accordingly for each sample. To compare accuracy, we define top-*k* accuracy as the percentage of the test samples in which the top *k*-ranked features belong to the set of relevant features. We implemented both methods for different sets of hyperparameters. We also compared consistency of the accuracy of these heuristic methods by repeating computations with 100 different random seeds for each set of hyperparameters.

#### 2.1.1 SensX achieves up to to 52% higher median accuracy

We computed the median accuracy across the 100 different runs for each set of hyperparameters (see Supplementary Figs 1 to 3). Table 2 compares the results for the best median accuracy achieved by the methods. SensX has higher median accuracy than SHAP in all cases, and also achieves a 100% accuracy in 8 cases whereas SHAP does so in 2 cases. Significant improvements are observed in nonlinear additive and switch data sets, in which SensX achieves up to 20% and 52% higher median accuracies than SHAP, respectively.

#### 2.1.2 SensX is up to 158 times faster in run-time

We compare the mean computation times for the 100 different runs of both methods with the sets of hyperparameters that gave the best median accuracy. Table 2 shows that SensX is up to 158 times and at least 21 times faster than SHAP for this analysis.

#### 2.1.3 SensX achieves higher consistency in all cases

We compare the consistency of the two methods by comparing the coefficients of quartile deviation (CQD) [Botta-Dukát, 2023] of accuracy from 100 runs with the sets of hyperparameters that gave the best median accuracy. A lower CQD implies higher consistency. Table 2 shows that SensX is more consistent than SHAP in all cases.

### 2.2 SensX validates vision transformer (ViT) models by explaining the characteristics of human facial attributes they have learned at full image resolution

Vision transformer (ViT) models [Dosovitskiy, 2020] apply the transformer architecture characterized by self-attention layers [Vaswani, 2017] to learn classification of images. We trained a ViT model on the publicly available Large-scale CelebFaces Attributes (CelebA) data set [Liu et al., 2015] to predict ‘Not Smiling’ and ‘Smiling’ for human faces. We denote this model by smiling-ViT. Similarly, we computed a ViT model that predicts eyeglasses on human faces, henceforth referred to as eyeglasses-ViT. Supplemental tables file shows that the labels for eyeglasses are severely imbalanced (6.5% true labels) whereas for smiling they are relatively balanced (48.2% true labels). However, both of the fine-tuned models show around a 0.91 F1-score on test data.

#### 2.2.1 SensX scales to more than hundreds of thousands of features

The input image has dimensions 224 *×* 224 pixels for each of the three color channels (RGB), totaling 150, 528 features. We were unable to use SHAP because it required at least 33 TB of memory to analyze a single test image, which was beyond the scope of the computational resources available to us. SensX takes at most 10 GB of memory and 7 minutes per sampled trajectory on a NVIDIA A100 GPU.

#### 2.2.2 SensX reveals relevant features at full resolution

Figure 3 shows the SensX values of features as heat maps, separately for the three color channels of a test image. It is visually evident that the pixels with high SensX values are around the mouth for the smiling-ViT. In contrast, pixels near the bridge of the nose and the frame of the eyeglasses have high SensX values with respect to the eyeglasses-ViT. Hence, our framework reveals that smiling-ViT and eyeglasses-ViT have learned intuitively correct facial attributes for smiling and eyeglasses, respectively, of this image.

**Figure 3:**
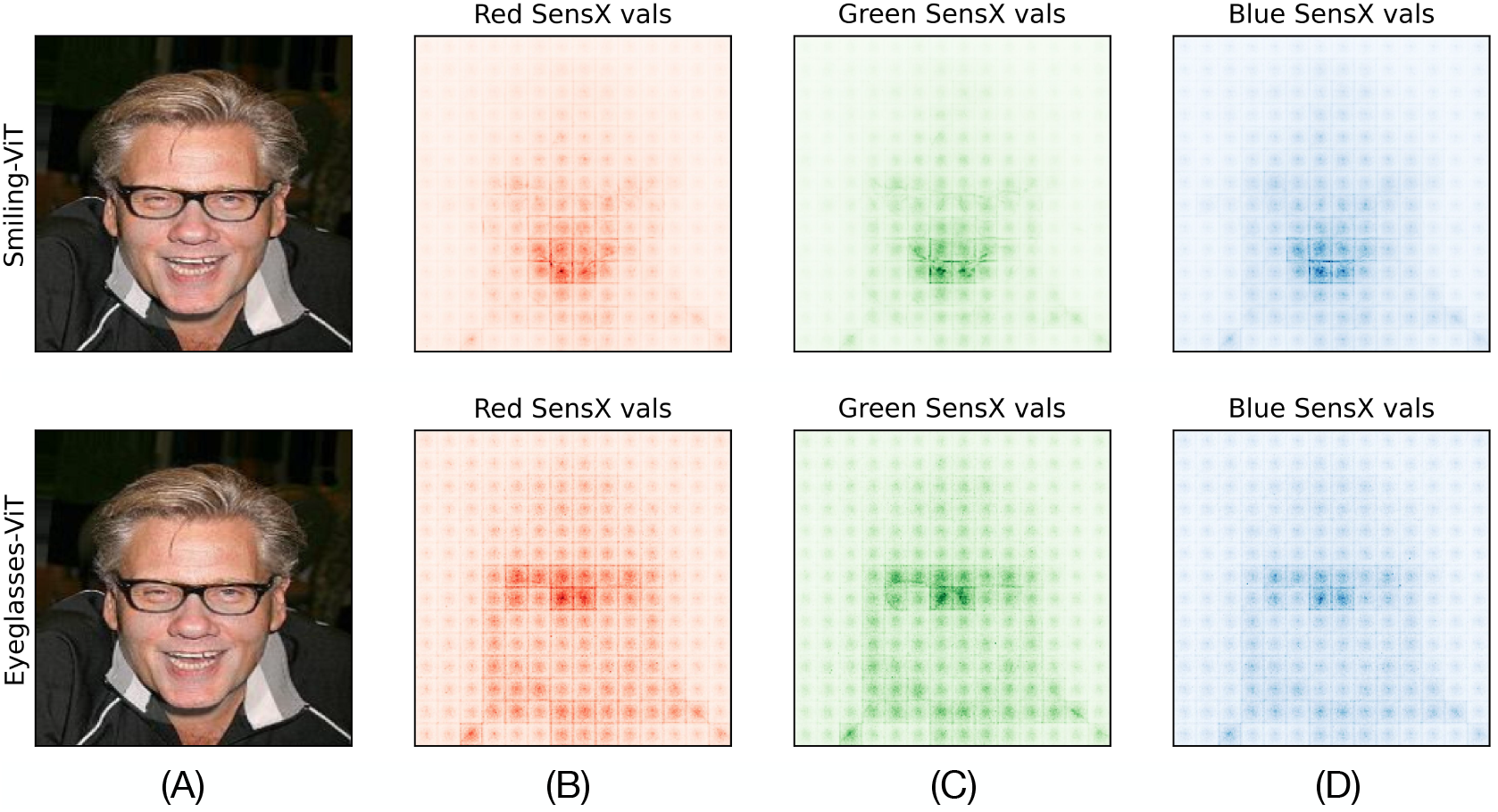
SensX validates that ViTs have learned inuitively accurate features as important for different facial attributes. (A) Test image. (B)-(D) Heatmaps of SensX values of pixels for red, green, and blue channels for smiling-ViT (top) and eyeglasses-ViT (bottom). Pixels around the mouth have high SensX values for the smiling-ViT, whereas pixels at the bridge of the nose and frame of the eyeglasses have high SensX values for the eyeglasses-ViT.

#### 2.2.3 Top ranked SensX features show an intuitively reasonable mask for smiling and eyeglasses learned by the ViT models

To see the mask of the test image defined by top *k* SensX features, we define a new image with top *k* SensX features set to their values in the test image and the remaining features set to 0. Figure 4 shows the resulting images for *k* = 1000, 2500, 5000, and 10000 for both smiling- and eyeglasses-ViT. It is visually evident that the top *k* SensX features of smiling-ViT define a mask of the test image around the mouth and capture the smile, while the top *k* SensX features of eyeglasses-ViT capture the bridge of the nose and the frame of the eyeglasses. Hence, the top SensX features create human interpretable masks of the test image that reasonably conform with what smiling- and eyeglasses-ViT should consider as important. *Can we determine an optimal value of k?* To answer this, we first introduce the SensX landscape.

**Figure 4:**
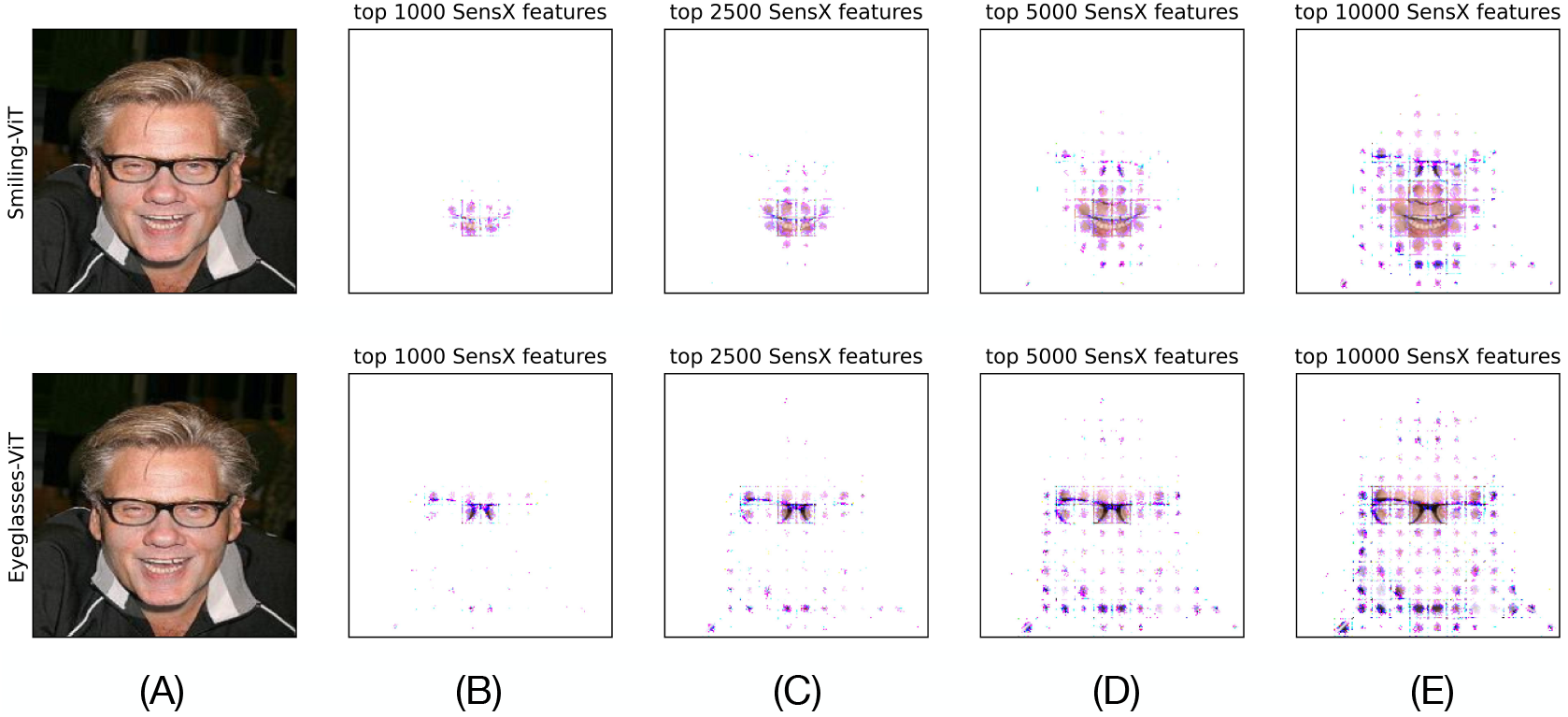
Top SensX features define intuitively accurate masks of different facial attributes.(A) Test image. (B)-(E) Images with top *k* = 1000, 2500, 5000, 10000 SensX features for smiling-ViT (top panel) and eyeglasses-ViT (bottom panel) having their corresponding value in the test image and the remaining features set to 0.

#### 2.2.4 SensX landscape defines multiple characteristics of the model

We conduct a more comprehensive analysis by perturbing the top *k* SensX features for a wider range of values of *k* and also for different values of the perturbation factor. The perturbation factor, denoted by *δ* ∈ [0, 1], defines the local neighborhood of the input sample to generate perturbations (see Methods). When *δ* = 0, the local neighborhood is the singleton set of the input sample. When *δ* = 1 it is the global domain. For each pair of values of *k* and *δ*, we compute the median of the model confidence for 1000 perturbations of the test image. The results are shown as interpolated heatmaps that we call SensX landscapes. Figure 5A shows SensX landscapes of the smiling- and eyeglasses-ViT for the test image. We make the following observations. (1) *δ* has to be greater than 0.2 and 0.5 to affect the median model confidence of the smiling- and eyeglasses-ViT models, respectively. (2) There exist distinct regions in the landscapes that result in a minimal value of the median model confidence. Importantly, the model confidence achieves its lowest value at *k* less than the total features. In other words, SensX landscape reveals a subset of features perturbing which reduces the model confidence lower than the model confidence when all features are perturbed.

**Figure 5:**
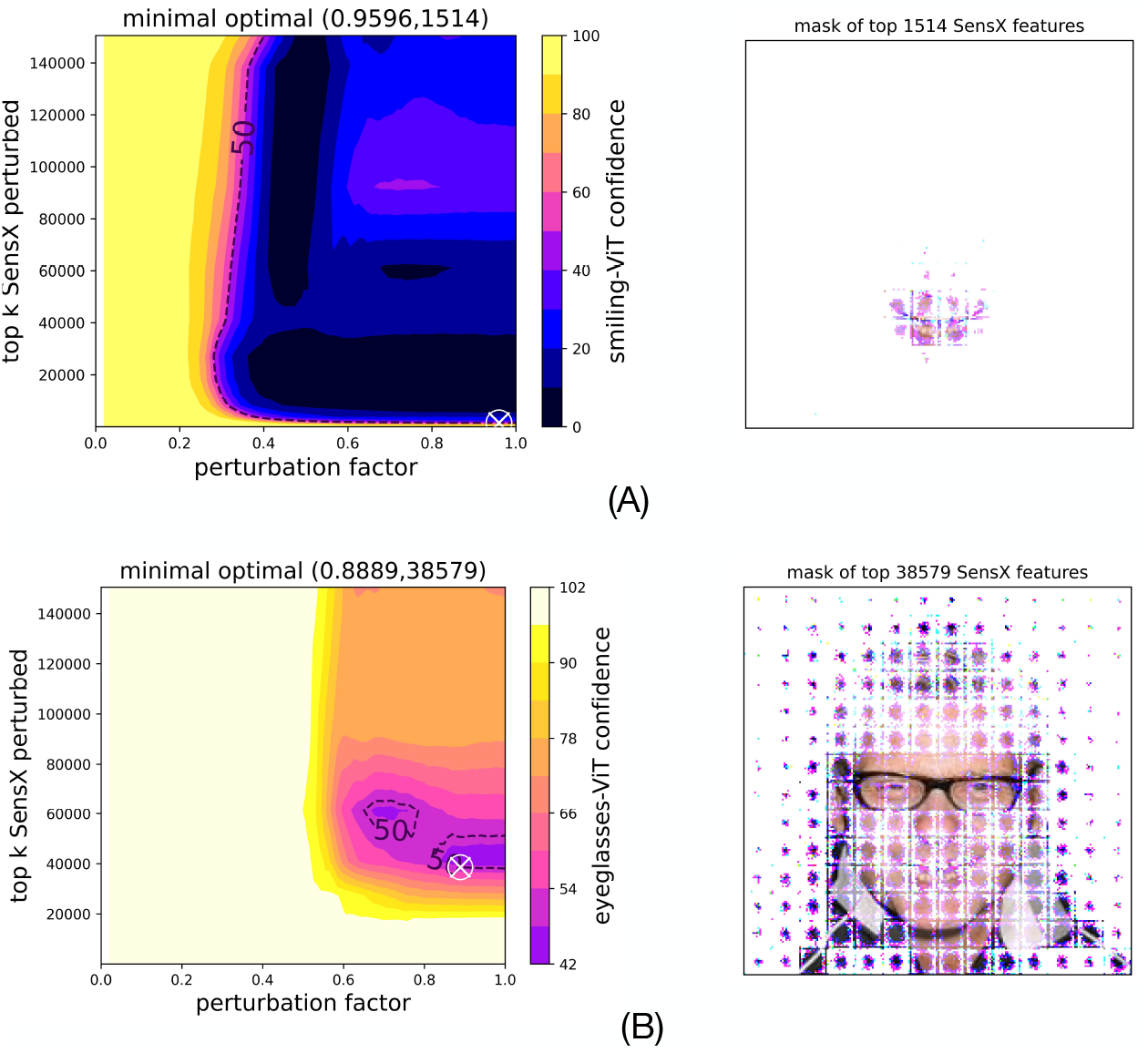
SensX landscapes give minimal sets of significant features. (A) Smiling-ViT and (B) Eyeglasses-ViT SensX landscapes (left panels) are heatmaps of median model confidence when different number of top SensX features are perturbed using different values of perturbation factors. ‘x’ marks a minimal set of features to be perturbed to reduce the model confidence below 50%. Right panels show the masks of the image defined by the minimal set of top SensX features.

#### 2.2.5 SensX landscape reveals a minimal set of only 1% of the total features that precisely demarcate the facial attributes that the smiling-ViT has learned as important

We define the minimal set of important SensX features as the smallest *k* such that perturbing top *k* SensX features reduces the model confidence below 50%. SensX landscape for smiling-ViT reveals that perturbing only 1514 features out of the total of 150, 528 features reduces the model confidence below 50%. Figure 5A (right panel) shows that the mask defined by these features demarcates the smile in the human face in the test image. On the other hand, perturbing at least top 38, 579 SensX features (26% of total features) reduces the median confidence of eyeglasses-ViT to below 50%. Figure 5B (right panel) shows the mask of these features in the test image.

#### 2.2.6 Instance-wise analysis shows that important features revealed by SensX are input-specific and accurate

Figure 6 shows the importance of input-wise XAI. SensX values show that the ViTs have correctly learned facial attributes specific to different human faces.

**Figure 6:**
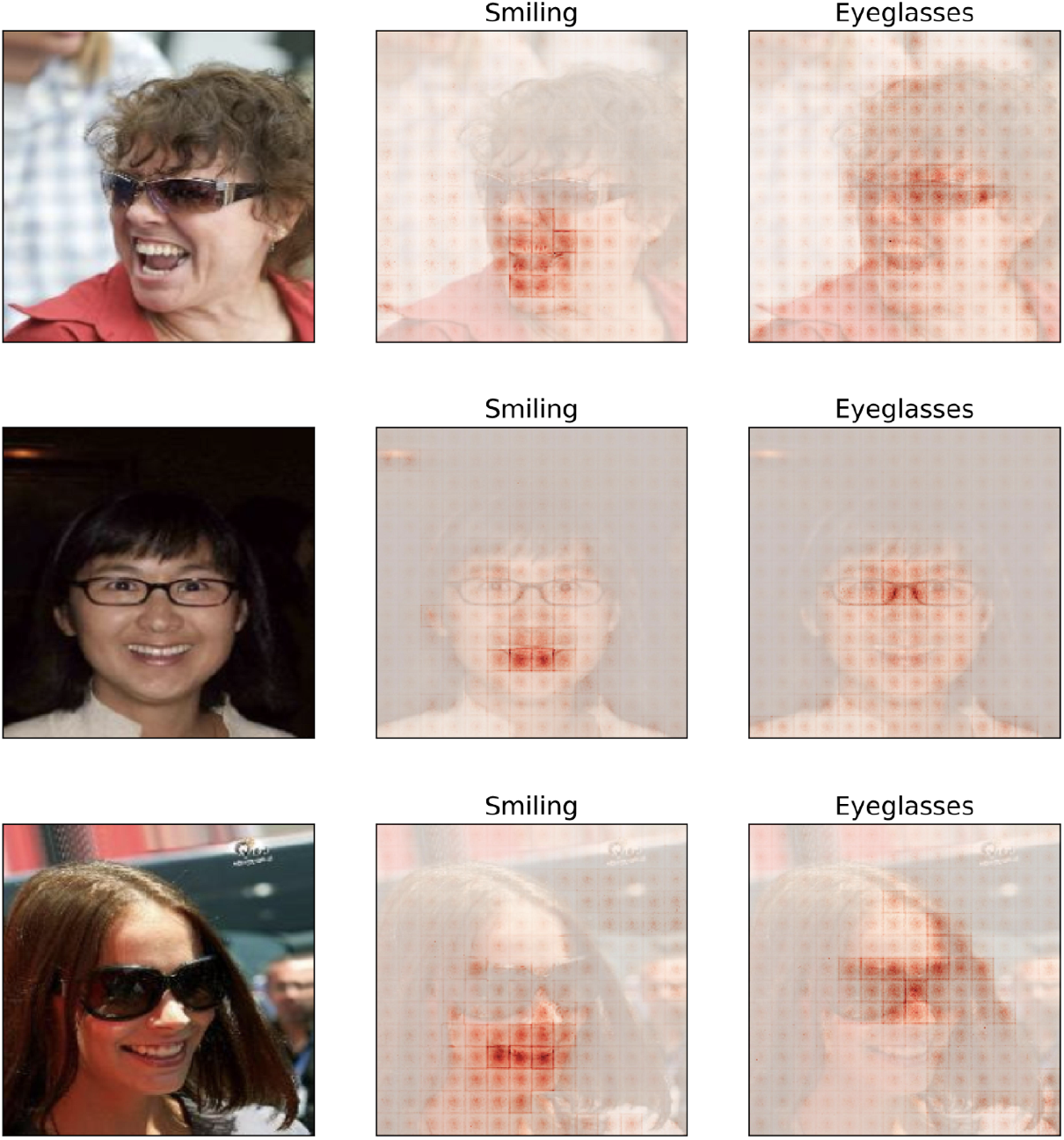
SensX validates that ViT models have learned input-specific important features accurately.

#### 2.2.7 SensX at full resolution reveals possible biases inherent to the model architecture

We notice that the heat maps of SensX values shown in Figure 3 reveal a grid-like structure—features on the grid boundaries and at the center of the grid patches are brighter than their neighbors. This grid is of size 16 × 16 that matches exactly with the patch-size of tokens in the ViT architecture [Dosovitskiy, 2020]. This suggests that there may be a bias towards the edges and center of the input tokens during training of this ViT architecture.

### 2.3 Automated cell type annotation using single-cell RNA data set

Single-cell RNA sequencing (scRNA-seq) quantifies gene expression levels in individual biological cells by measuring abundance of mRNA molecules per cell [Heumos et al., 2023]. Advancements in experimental technology have resulted in large data sets with tens of thousands of genes measured in millions of individual cells. Annotating cell types using scRNA-seq data is a fundamental problem [Hou and Ji, 2024]. Various automated methods for cell type annotation using different AI architectures have been proposed, for example, transformers [Chen et al., 2023], graph neural networks [Shao et al., 2021], and FNNs [Ma and Pellegrini, 2020]. For biological validation and hypotheses development, it is necessary to explain what black-box AI models trained for automated cell annotation have learned as distinguishing genes for different cell types.

We implemented FNNs to learn classification of cell types using the publicly available scRNA-seq data set of the human lung cell atlas (HLCA) [Sikkema et al., 2023]. The data set has 1, 305, 099 cells with gene expression values reported for 56, 239 genes per cell. We trained binary classifiers to identify 21 different cell types (see Figure7 for a list of the cell types).

#### 2.3.1 SensX vectors cluster distinctly for different cell types

We computed SensX values for 1000 cells of each cell type from the test data for which the model predicted with at least 99% confidence that they belonged to their respective class. A three-dimensional embedding of the gene expression values of these cells does not show distinct clusters for the different cell types (see Figure 7A). However, Figure 7B shows an embedding based on the SensX distance between cells (see Methods 4.1.6 for definition) that visually suggests different clusters. This suggests that the DNNs trained to identify different cell types have learned different sets of genes as important.

**Figure 7:**
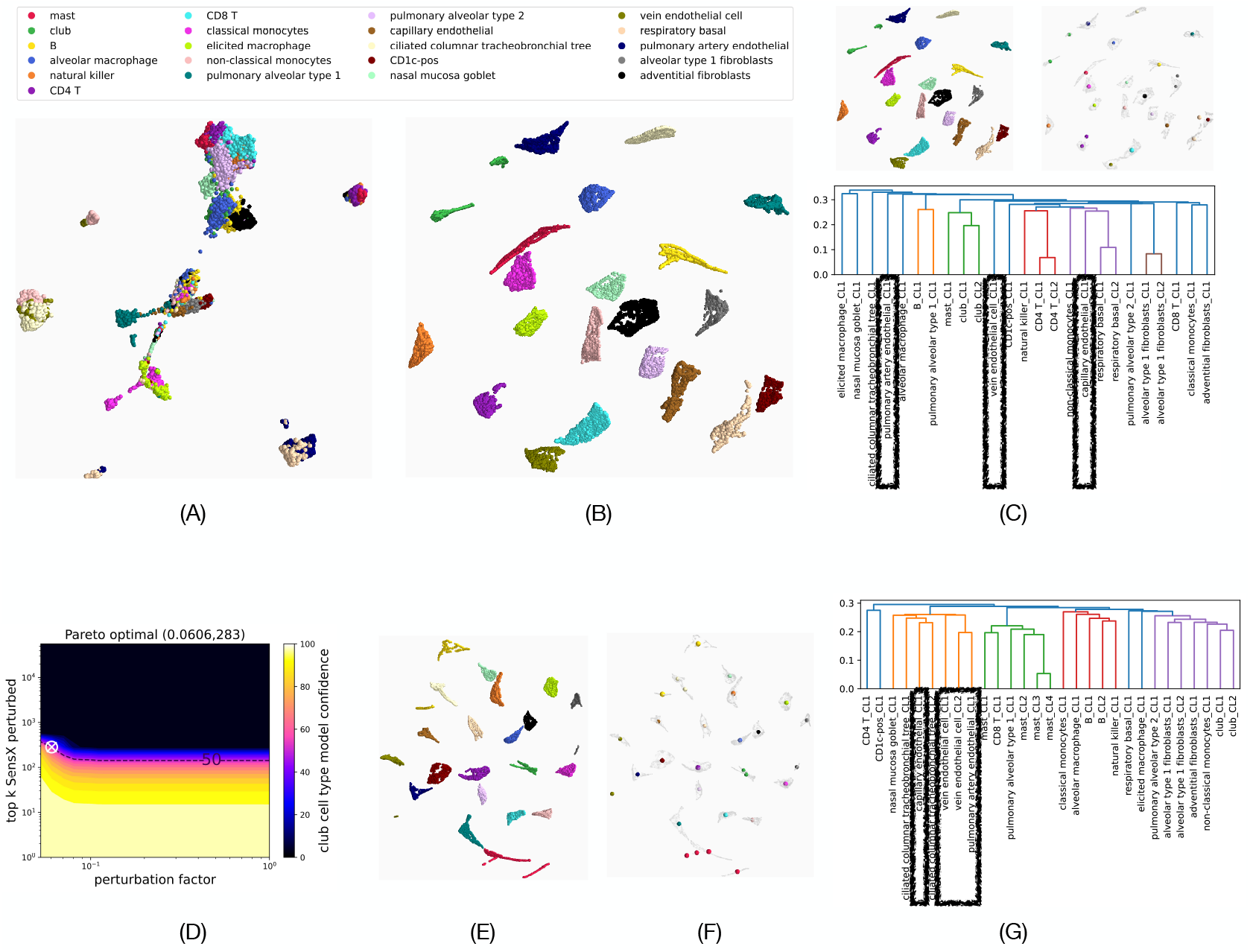
Optimal set of important SensX features contain biologically significant information. (A) UMAP [McInnes et al., 2018] 3D-embedding of the log-normalized gene expression values of 1000 cells of each cell type. (B and C top left panel) UMAP 3D-embedding based on pairwise SensX distances between cells.(C top right panel) Means of DBSCAN clusters (C bottom panel) Dendrogram of means of DBSCAN-clusters to show heirarchical clustering. (D) SensX landscape of a single mast cell. ‘x’ marks the Pareto optimal that minimizes the number of top SensX features to be perturbed and the perturbation factor. (E) UMAP 3D embedding of the SensX set distances of the sets of significant features of each cell. (F) Means of clusters identified by DBSCAN. (G) Dendrogram of hierarchical clustering of the means of DBSCAN-clusters.

For a more rigorous clustering analysis than visual inspection, we used DBSCAN [Ester et al., 1996] to cluster the embedding of the SensX distances (Figure 7B and also Figure 7C top-left panel). DBSCAN determined 25 clusters and 0 outliers. Each cluster contained cells of a single cell type, confirming that there are distinct clusters for the different cell types. For some cell types, DBSCAN identified multiple clusters. We computed the mean of each cluster (see Figure 7C top-right panel) and labeled the mean of the cluster number *k* of cell type ct by ⟨ct⟩ _CL*k*. Finally, we performed hierarchical clustering [Müllner, 2011] of the means and show the resulting dendrogram in Figure 7C bottom panel. We observe that different DBSCAN-clusters of the same cell types (club, CD4T, respiratory basal, alveolar type1 fibroblasts) are close to each other in the dendrogram, providing evidence that SensX ranking is biologically meaningful. However, the clusters of three different endothelial cell types (pulmonary artery, vein, capillary) are far from each other in the dendrogram-tree. *We claim that this is due to mapping insignificant SensX values to integer ranks*. To prove this, we first define and determine a set of genes with significant SensX ranks for each cell.

#### 2.3.2 SensX determines that only 0.5% of total genes are important genes for most cell types

To compute an optimal set of important genes, we consider both the number of top *k* genes and *δ*. This is because a higher value of *δ* might result in a local neighborhood of the input that spans multiple log scales because the gene expression values in scRNA-seq data are log-normalized. Hence, it may be of biological interest to minimize *δ* as well to avoid perturbations that imply changing gene expression values too widely. Hence, we define the Pareto optimal as the pair (*k, δ*) with minimal norm that reduces the median model confidence below 50%. Figure 7D shows SensX landscape of a single mast cell. The Pareto optimal point is marked by an ‘x’. Supplemental tables file shows the median and CQD values of the Pareto optimal points computed for 1000 cells of each cell type. For all but one cell type (mast), perturbing 284 top SensX genes (0.5% of the total genes) by a perturbation factor of 0.1 to 0.25 reduces the median model confidence below 50%.

#### 2.3.3 Sets of significant SensX features reveal biologically relevant relations between different cell types

Using the computed optimal sets of features for each cell, we compute the SensX set distance (see Methods 4.1.7) between every pair of cells. Briefly, SensX set distance measures set dissimilarity between two sets of significant features, weighted by the discrepancy in the ranks of the features between the two sets. Figure 7E shows a three-dimensional embedding based on the pairwise SensX set distances. Figure 7F shows the means of 29 clusters that are identified by DBSCAN, showing that the significantly ranked features are different for different cell types. Figure 7G shows the dendrogram from hierarchical clustering of the means of DBSCAN-clusters, revealing that indeed the sets of significantly ranked features of the three different endothelial cell types are highly similar.

## 3 Discussion

We introduced the notion of justifiable perturbations to systematically conduct GSA. We designed SensX, a model agnostic framework for XAI that uses justifiable perturbations and GSA to explain black-box DNNs. Benchmarks using synthetic data sets showed that SensX outperformed the current most widely used XAI method in accuracy, computation time, and consistency. SensX was the only XAI that scaled to analyze ViT models that have more than 150, 000 features. It validated the ViT models trained to identify smiling and eyeglasses and revealed biases inherent to the model architecture. We used SensX to explain DNNs trained to annotate cell type using scRNA-seq data. It determined subsets of genes important to different cell types, paving way for hypotheses development and further experimental design.

We showed how SensX can be used to determine optimal subsets of important features which had as few as 0.5% of the total number of features in some cases. This significantly reduces the dimensionality of any further analyses that may be used to determine precise correlations between the important features.

SensX is not limited to supervised classification learning and generalizes to regression problems by choosing the quantity of interest accordingly. We analyzed multiclass data sets using binary classifiers in two different ways. First, we used used transfer learning to fine-tune a pretrained ViT multiclass classifier to a binary classifier. Second, we used one-vs-all binary classifiers to learn annotation of multiple cell types using scRNA-seq data set.

The underlying assumption in the framework is that uniformly distributed perturbations significantly affect the QOI. Therefore, the QOI should be chosen judiciously. SensX provides a criterion to justify the choice of QOI—*δ*^*^ *>* 0 (see Methods). The QOI of the case studies was chosen as the confidence of the model that the given input belongs to the class. It is generally assumed that uniform perturbations represent noise and that data representing a specific class is distinguished by a specific pattern or relations in the features that are not uniformly distributed. However, this may not always be the case (see Supplementary Figure 5).

The number of trajectories is an important hyperparameter of SensX to balance the convergence of GSA measures and computation time. There is no theoretical basis to compute an optimal value, but we provide an empirical strategy to make a judicious choice. A computational advantage of SensX is that it can be computed in batches without any additional overhead, and the results of any number of batches can be combined. Hence, it is advisable to perform computations in batches, and the convergence of GSA measures can be assessed by combining all batches (see Supplementary Figure 4).

We note that all perturbations are uniform and independent within the justifiable domain. We do not assume any correlations between the features a priori. The aim of SensX is to determine the correlations between features that a trained model has learned, and it does so by GSA using independent perturbations to explore the input space. Given the trained model, the optimal set of top SensX features corresponds to an optimal set of correlated features that may be further analyzed to determine precise signed correlations.

SensX presents a general model agnostic framework to decide the domain of perturbations for GSA to explain DNNs. The domain of perturbations for perturbations-based GSA in any context should be specified with care. Model-specific methods to reasonably determine the domain according to the underlying process being studied have been previously suggested [Aggarwal et al., 2024, 2019, 2018].

## 4 Methods

### 4.1 Framework of SensX

#### 4.1.1 Terminology

A trained model parameterized by a fixed set of parameters is a mathematical function that maps an input to an output, *m* : ℝ^*n*^ →ℝ^1^. **x** = (*x*_1_, …, ℝ_*n*_) ∈ ∝^*n*^ is a model input of *n* features and **y** = *m*(**x**) ∈ ℝ^*l*^ is the corresponding model output. The quantity of interest for GSA is a function of the model input,

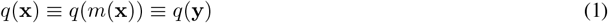

with range 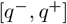.

#### 4.1.2 Meaningful perturbations: Global domain and local neighborhoods

We define the global domain as

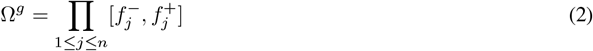

where 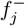 and 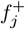 are lower and upper bounds of the *j*-th feature. The framework accepts any user-defined Ω^*g*^ that may be contextually relevant based on expert advice. In each application shown in this work, Ω^*g*^ was chosen as the smallest hypercube containing the training set.

We define a local neighborhood of **x** ∈ Ω^*g*^ parameterized by perturbation factor *δ* ∈ [0, 1] as

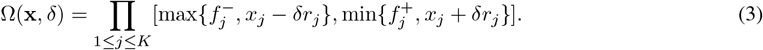

From equation 3

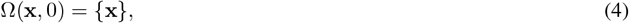

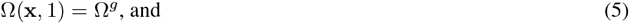

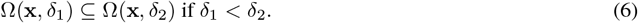

#### 4.1.3 Significant perturbations: Optimal local neighborhood

We introduce four hyperparameters,

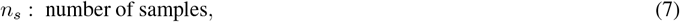

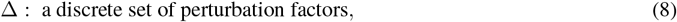

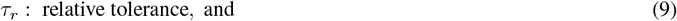

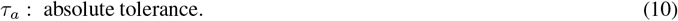

The idea is to find the maximal perturbation factor *δ*^*^ ∈ [0, 1] such that the median of QOI of *n*_*s*_ perturbations in Ω(**x**, *δ*) does not change significantly ∀*δ* ∈ (*δ*^*^, 1].

We denote a set of *n*_*s*_ perturbations in Ω(**x**, *δ*) by

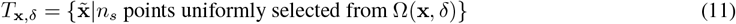

and the corresponding median of the quantity of interest of perturbations by

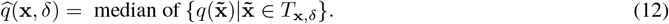

We define optimal perturbation factor 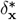 as the maximal value of *δ* ∈ Δ with 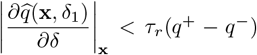 and 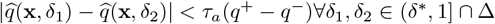.The optimal local neighborhood is defined by 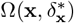. If 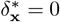, then SensX is not valid because the domain of perturbations will be the trivial set Ω(**x**, 0) = {**x**}.

By default, *n*_*s*_ = 1000, *τ*_*r*_ = *τ*_*a*_ = 0.1, and Δ = linspace(0.02, 1, num = 50).

#### 4.1.4 Grounded trajectories: Modified Morris method to compute SensX values

Let 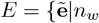 samples uniformly selected from 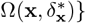 where *n*_*w*_ is a hyperparameter that denotes the number of walks. We define a trajectory in 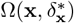 starting at **x** and ending at some 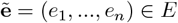 as a tuple of points

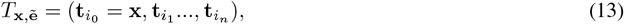

where (*i*_1_, …, *i*_*n*_) is a randomly permuted sequence of integers from 1 to *n*, and 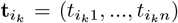 such that for *k >* 0

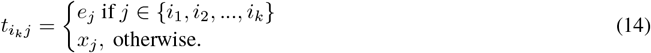

We call 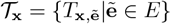 a set of trajectories grounded in **x**.

#### 4.1.5 SensX values and SensX vector

The elementary effect of feature *i*_*k*_ along a trajectory 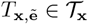 is defined by

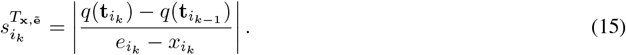

We define the SensX value of feature *i*_*k*_ with respect to an input **x** as the average of elementary effects over a set of grounded trajectories,

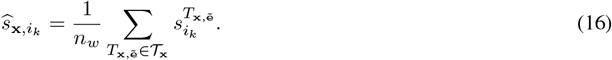

Finally, we define the SensX vector with respect to input **x** as

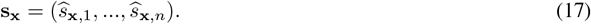

#### 4.1.6 SensX distance: Comparing SensX values of multiple inputs

SensX vectors of two inputs cannot be compared directly because SensX values are not normalized measures. Therefore, we instead compare the ranks of SensX features as follows. We define the SensX rank vector as

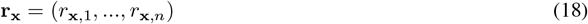

where *r*_**x**,*k*_ is the rank of 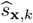 in **s**_**x**_.

We define the SensX distance between two inputs **x**_1_, **x**_2_ ∈ Ω^*g*^ as

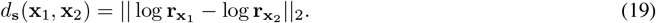

#### 4.1.7 SensX set distance: Comparing sets of important features of multiple inputs

The idea is to measure a dissimilarity of the sets of important features that is also weighted by the discrepancies in the ranks of the features.

Let *F* and *G* be the sets of important features of inputs **x**_1_ and **x**_2_, respectively. Let *H* = *F* ∪*G* = {*h*_1_, …, *h*_*m*_}. We define two boolean vectors

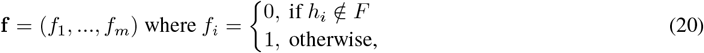

and

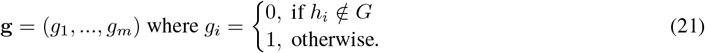

We define SensX set distance between inputs **x**_1_ and **x**_2_ as the weighted Jaccard dissimarity Kaufman and Rousseeuw [2009] between **f** and **g**,

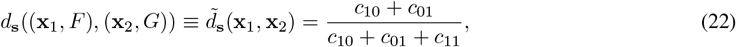

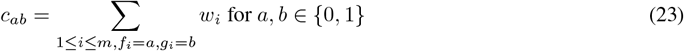

where we define

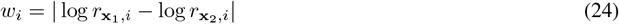

as the weight of feature *h*_*i*_ ∈ *H* that accounts for the discrepancy in ranks of feature *i*.

### 4.2 Computational implementation

We implemented SensX in Python using the Google Just After eXecution (JAX) machine learning library [Bradbury et al., 2018]. For computational efficiency, we designed our algorithm using pure functions and without side-effects to allow just-in-time (JIT) compilation. We implemented the deep learning models in this work using NNX, a JAX-based neural network library. Supplementary file `conda_packages.txt` shows versions of all packages in the conda environment to reproduce the results. Model simulations were done on NVIDIA A100 GPUs. Supplemental table file shows the values of SensX hyperparameters used for the models in this work.

For efficient parallelization, two different algorithms to compute SensX are provided—batching over inputs or data samples to be analyzed and batching over the features per data sample. The latter is recommended if the number of inputs is very few as compared to the number of features. Since a trajectory has an inherent order, batching over features using pure functions that allow JIT compilation is non-trivial.

### 4.3 Synthetic data sets

#### 4.3.1 Generating labeled data

Samples are generated from ten-dimensional standard Gaussian distribution. Each sample, **x** = (*x*_1_, …, *x*_10_) is labeled *b* ∈ {0, 1} based on the following criteria for three of the four synthetic data sets (https://github.com/Jianbo-Lab/CCM>).

- XOR model: *P* (*b* = 1) ∝ *x*_1_*x*_2_
- orange skin model: 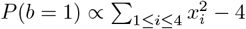
- nonlinear additive model: 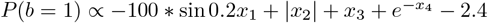

The switch data set is a mixture of data samples from two different criteria as follows. *x*_10_ is generated from a mixture of two Gaussians centered at ±3 with equal probability. If *x*_10_ is generated from the Gaussian centered at 3, then *x*_1_ to *x*_4_ are used to label the sample based on the orange skin model. Otherwise, *x*_5_ to *x*_8_ are used to label the sample based on the nonlinear additive model.

#### 4.3.2 Model and training

We generated 100, 000 labeled data samples for each data set and used FNNs with layers [10, 200, 100, 50, 2] to learn the classifications. We used 80% of the samples as the training set and the rest as the test set. We trained the models by minimizing the crossentropy loss using the AdamW method with the constant learning rate of 0.0001. Each optimization step consisted of a batch of 100 randomly-picked samples from the training set. We implemented 10, 000 optimization steps and computed the F1-score for the test set after every 100 steps. The models analyzed using SHAP and SensX were defined by the sets of parameters that yielded the maximal F1-score (see Supplemental file `supplemental_tables.xlsx` for the scores).

#### 4.3.3 Multiple sets of hyperparameters for XAI methods

SHAP: Number of samples *n*_*s*_ = {100, 500, 1000}

SensX: *n*_*w*_ = {100, 500, 1000, 2500, 5000, 10000, 15000}

### 4.4 Vision transformer (ViT)

#### 4.4.1 Model setup

We implemented the vision transformer model defined at https://docs.jaxstack.ai/en/latest/JAX_Vision_transformer.html and loaded the model parameters pre-trained on the ImageNet-21k data set from the transformers Python library [Wolf et al., 2020]. We replaced the model classifier with a fully connected layer with two outputs.

To compute smiling-ViT, we fine-tuned the model by training it on the publicly available images of human faces from the Large-scale CelebFaces Attributes (CelebA) data set [Liu et al., 2015] to predict label 0 for faces annotated not smiling and to predict label 1 for those annotated smiling. Similarly, we computed eyeglasses-ViT to predict eyeglasses on image of a human face.

#### 4.4.2 Fine-tuning to Smiling and Eyeglasses

CelebA data set contains 202, 599 images. Train:test split is chosen as 80 : 20. We trained the models by minimizing the crossentropy loss using the SGD method with a constant learning rate of 0.00001 and Nesterov momentum of 0.8. Each optimization step consisted of a batch of *n* = 25 randomly-picked samples from the training set. For balanced training, we ensured that ⌊*n/*2⌋ of the samples belonged to the class and the rest did not. We implemented 60, 000 optimization steps and computed the F1-score for the test set after every 1000 steps. The models analyzed using SHAP and SensX were defined by the sets of parameters that yielded the maximal F1-score (see Supplemental file `supplemental_tables.xlsx` for the scores).

The five example images shown are randomly selected from images for which both smiling-ViT and eyeglasses-ViT had at least 99% model confidence.

### 4.5 Single-cell data set

#### 4.5.1 Model training

We used FNNs with layers [56239, 250, 50, 2] to learn the annotations with different cell types. Train:test split is chosen as 80 : 20. We generated the train and test sets such that both have approximately the same percentage composition of the different cell types. Each optimization step consisted of a batch of 500 randomly-picked samples from the training set. For balanced training, we ensured that the percentage composition of the batch matched the percentage composition of the data set. We implemented 7, 500 optimization steps and computed the F1-score for the test set after every 500 steps. The models analyzed by SensX were defined by the sets of parameters that yielded the maximal F1-score (see Supplemental file supplemental_tables.xlsx for the scores).

### 4.6 Code availability

Computer code to reproduce results is available at https://github.com/nihcompmed/SensX.

## Acknowledgements

This research was supported by the Intramural Research Program of the NIH, the National Institute of Diabetes and Digestive and Kidney Diseases (NIDDK), project number Z01-DK075146. This work utilized the computational resources of the NIH HPC Biowulf cluster (https://hpc.nih.gov).

