## Supplemental figures for "Sensitivity based model agnostic scalable explanations of deep learning"

Supplementary information

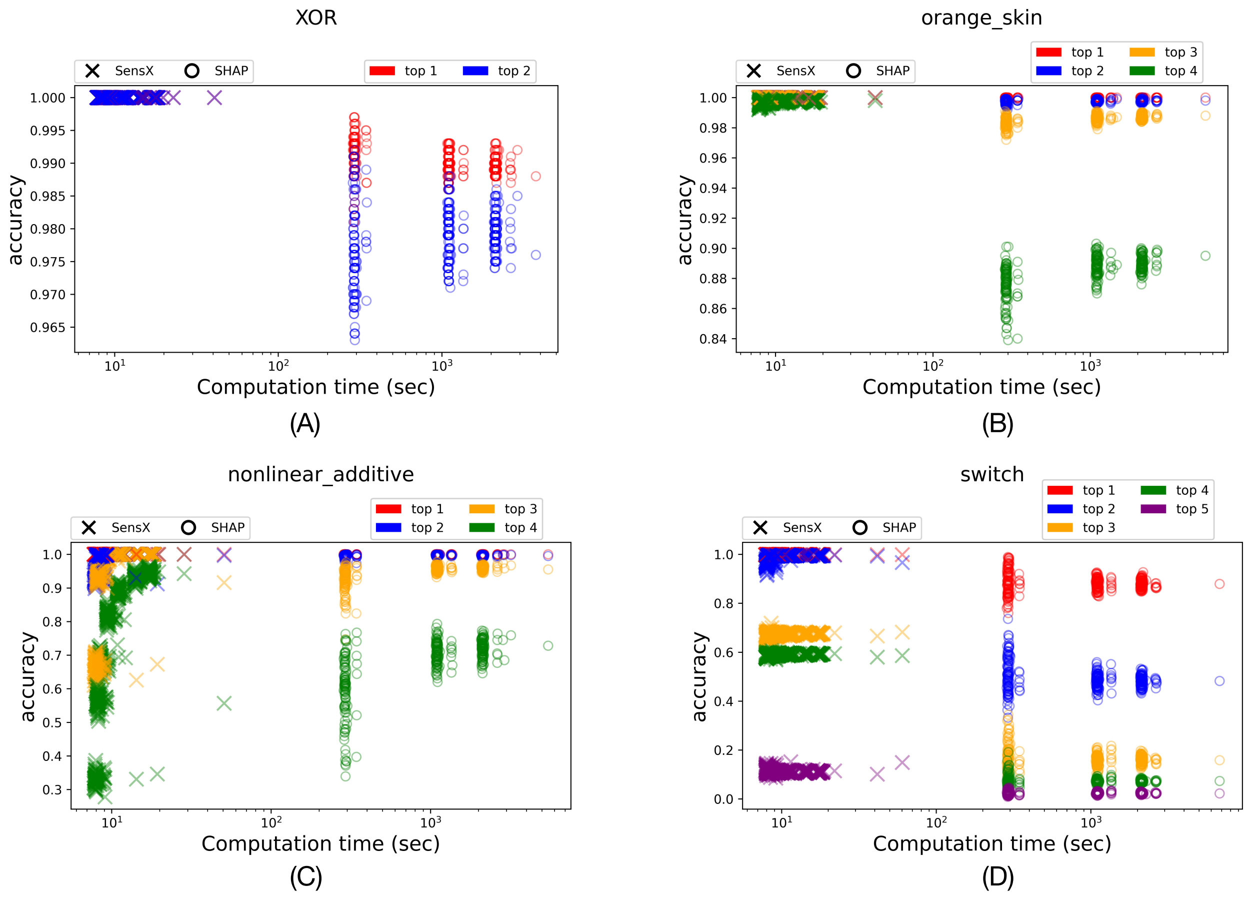


Fig 1: Benchmarks for all 100 runs for all sets of parameters for SHAP and SensX for synthetic data sets (A) XOR (B) orange_skin (C) nonlinear_additive and (D) switch.

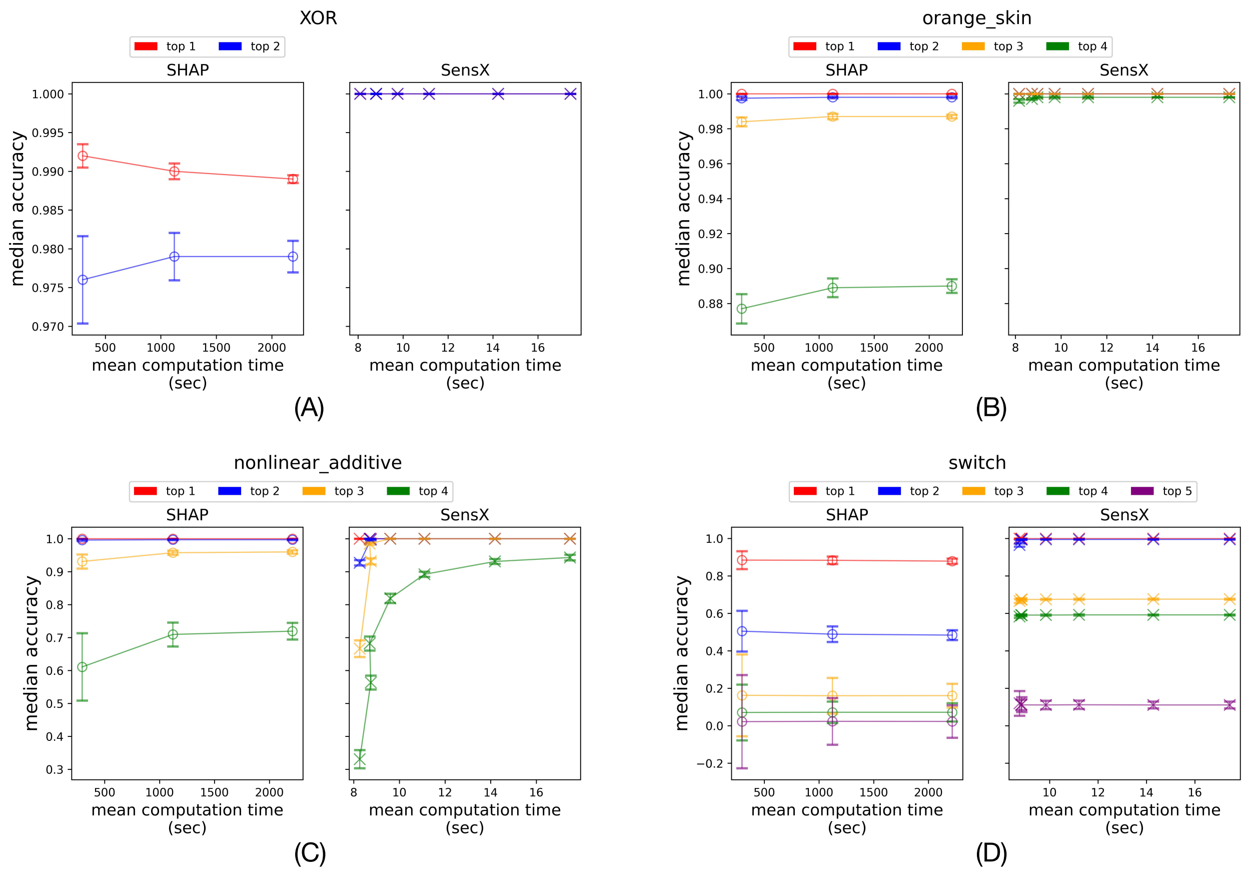


Fig 2: Mean time taken vs. median accuracy for SHAP and SensX for synthetic data sets (A) XOR (B) orange_skin (C) nonlinear_additive and (D) switch.

.

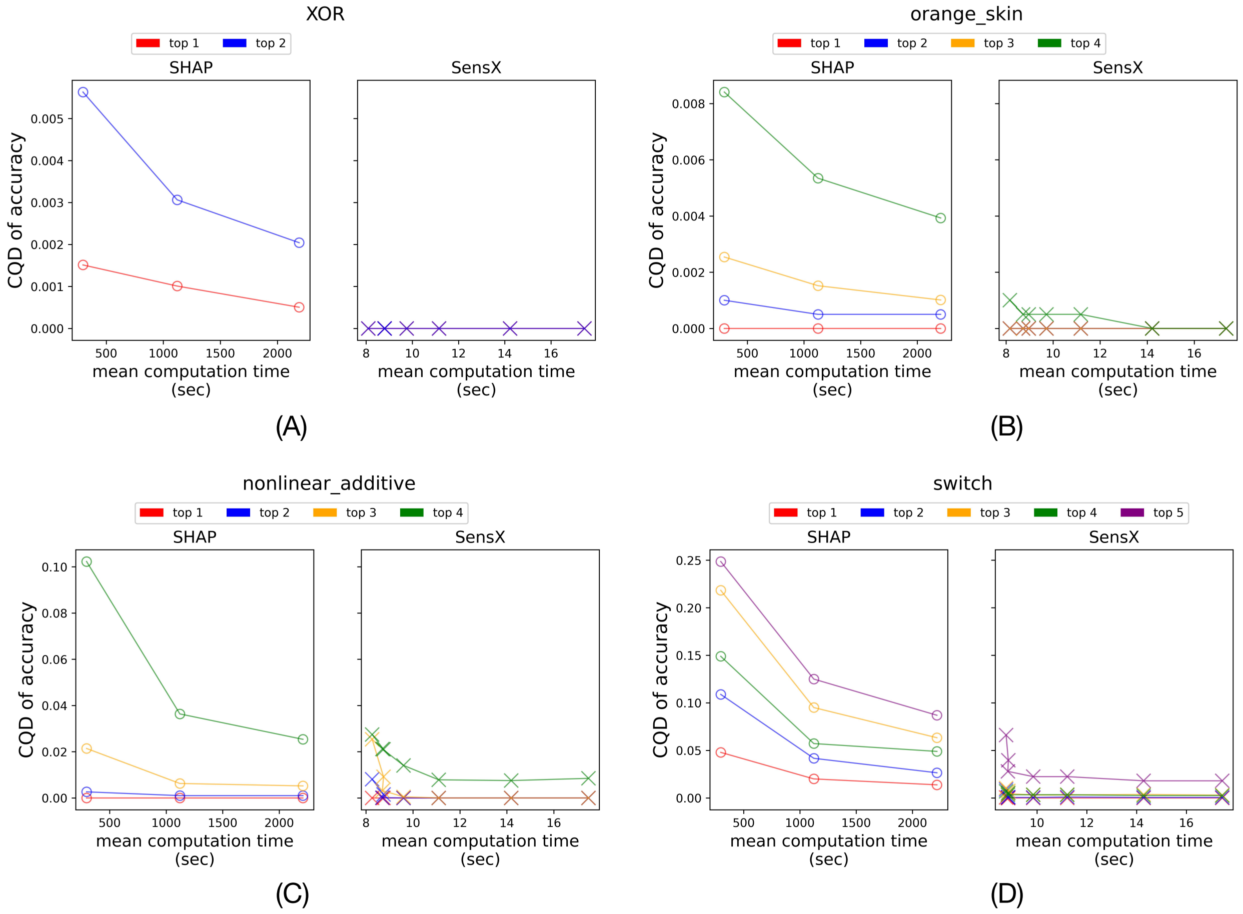


Fig 3: Coefficient of quartile deviation (CQD) in accuracy vs. mean time taken for SHAP and SensX for synthetic data sets (A) XOR (B) orange_skin (C) nonlinear_additive and (D) switch.

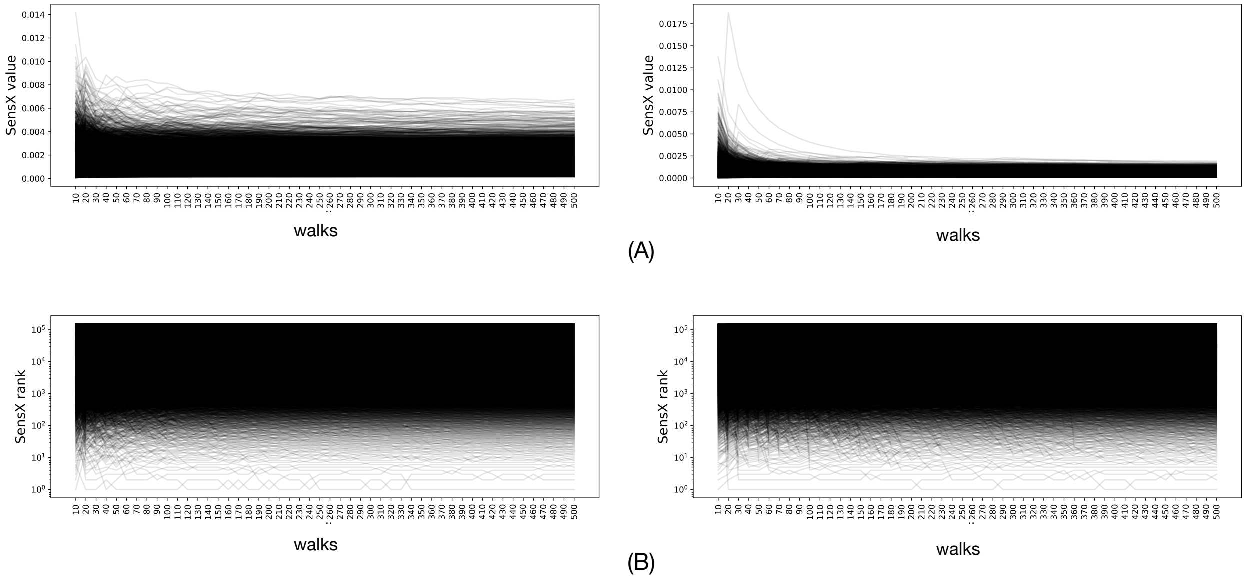


Fig 4: (A) SensX values and (B) SensX ranks converge as the number of walks is increased.

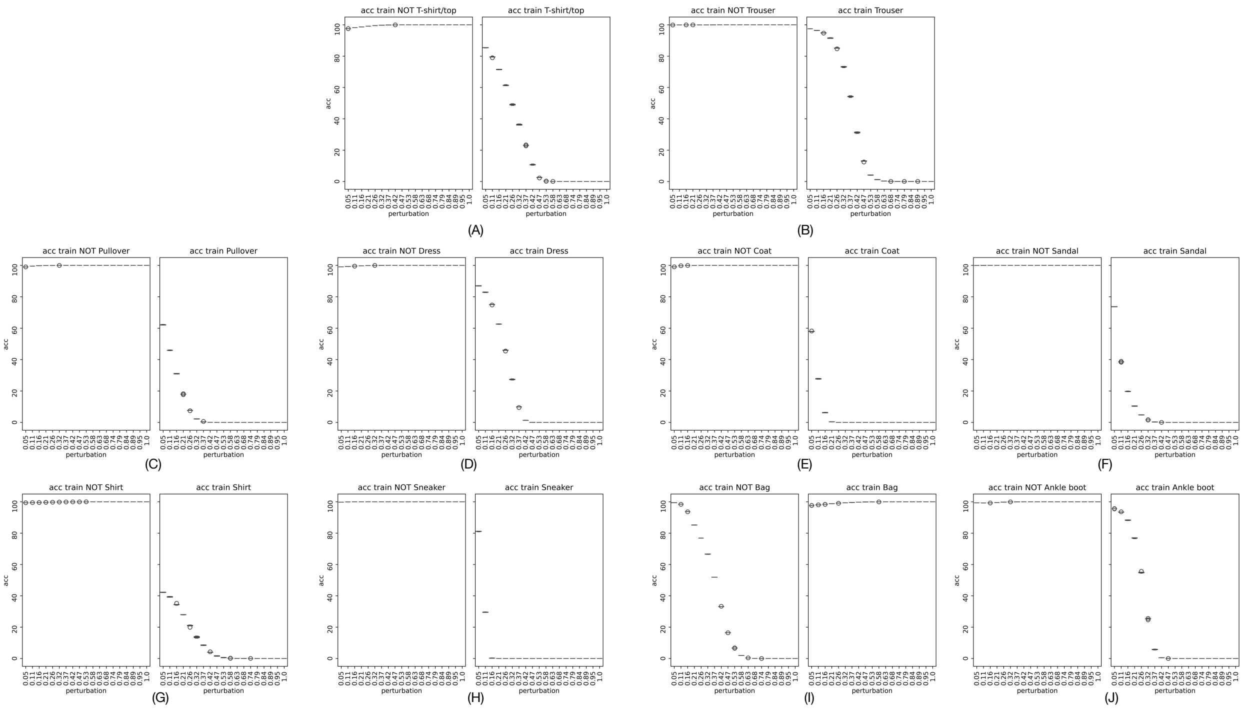


Fig 5: We trained FNNs one-vs-all binary classifiers to learn Fashion MNIST classification. Perturbations reduce the accuracy in identifying label 1 in all cases but one. Uniform perturbations in inputs to the (I) bag-classifier reduce the accuracy in identifying label 0 instead of label 1 because the true pattern for a bag is a uniform distribution on an average and every other class has a non-uniform distribution.
