## Supplemental tables for "Sensitivity based model agnostic scalable explanations of deep learning"

| Model/datase | F1-score on test data |
|---|---|
| XOR | 0.99 |
| orange_skin | 0.97 |
| nonlinear_a | 0.99 |
| switch | 0.98 |

| model | top-k | median accuracy (%) | | accuracy differential (%) | mean time taken (sec) | | speed-up factor | CQD (lower is better) | |
|---|---|---|---|---|---|---|---|---|---|
| | | SHAP | SensX | | SHAP | SensX | | SHAP | SensX |
| XOR | 1 | 99.2 | 100 | **+0.8** | 296 | 14 | **21** | 0.0015 | **0** |
| | 2 | 97.9 | 100 | **+2.1** | 2190 | 14 | **156** | 0.002 | **0** |
| orange_skin | 1 | 100 | 100 | | 2206 | 14 | **158** | 0 | 0 |
| | 2 | 99.8 | 100 | **+0.2** | 2206 | 14 | **158** | 0.0005 | **0** |
| | 3 | 98.7 | 100 | **+1.3** | 2206 | 14 | **158** | 0.001 | **0** |
| | 4 | 89 | 99.8 | **+10.8** | 2206 | 14 | **158** | 0.0039 | **0** |
| nonlinear_additive | 1 | 100 | 100 | | 2210 | 14 | **158** | 0 | 0 |
| | 2 | 99.7 | 100 | **+0.3** | 2210 | 14 | **158** | 0.001 | **0** |
| | 3 | 96 | 100 | **+4.0** | 2210 | 14 | **158** | 0.0052 | **0** |
| | 4 | 72 | 94.3 | **+22.3** | 2210 | 17 | **130** | 0.0254 | **0.0085** |
| switch | 1 | 88.5 | 99.9 | **+11.5** | 296 | 14 | **21** | 0.0479 | **0** |
| | 2 | 50.5 | 99.5 | **+49.0** | 296 | 14 | **21** | 0.1089 | **0.0005** |
| | 3 | 16.3 | 67.6 | **+51.3** | 296 | 14 | **21** | 0.2183 | **0.0037** |
| | 4 | 7.2 | 59.2 | **+52.0** | 2218 | 14 | **158** | 0.049 | **0.0025** |
| | 5 | 2.4 | 12 | **+9.6** | 1123 | 9 | **125** | 0.125 | **0.0661** |

**Eyeglasses-ViT**

| | Train | Test | Total |
|---|---|---|---|
| FALSE | 151525 | 37881 | 189406 |
| TRUE | 10555 | 2638 | 13193 |
| % TRUE | 6.51221619 | 6.51052593 | 6.51187814 |

**Smiling-ViT**

| | Train | Test | Total |
|---|---|---|---|
| FALSE | 83944 | 20986 | 104930 |
| TRUE | 78136 | 19533 | 97669 |
| % TRUE | 48.2082922 | 48.207014 | 48.2080366 |

| Models | F1-score on test |
|---|---|
| Smiling | 0.91 |
| Eyeglasses | 0.92 |

| cell type | F1 score test data |
|---|---|
| lveolar macrophage | 0.97572337 |
| natural killer | 0.9717783 |
| CD4 T | 0.93786632 |
| CD8 T | 0.93556662 |
| Classical monocytes | 0.94307875 |
| elicited macrophage | 0.89077587 |
| Non-classical monocytes | 0.92085559 |
| pulmonary alveolar type 1 | 0.97926925 |
| pulmonary alveolar type 2 | 0.99042482 |
| capillary endothelial | 0.97593563 |
| ciliated columnar tracheobronchial tree | 0.99126953 |
| CD1c-pos | 0.91643286 |
| nasal mucosa goblet | 0.95301045 |
| vein endothelial cell | 0.96219065 |
| respiratory basal | 0.98048556 |
| mast | 0.99329033 |
| pulmonary artery endothelial | 0.94153758 |
| alveolar type 1 fibroblasts | 0.98639859 |
| adventitial fibroblasts | 0.97847957 |
| club | 0.92358233 |
| B | 0.99393129 |

| cell type | perturbation factor | | top k SensX perturbed | |
|---|---|---|---|---|
| | median | CQD | median | CQD |
| mast | 0.24 | 0.14 | 566 | 0.33 |
| club | 0.09 | 0.18 | 284 | 0 |
| B | 0.25 | 0.14 | 284 | 0.33 |
| alveolar macrophage | 0.14 | 0.17 | 284 | 0 |
| natural killer | 0.14 | 0.24 | 284 | 0 |
| CD4 T | 0.09 | 0.26 | 284 | 0 |
| CD8 T | 0.12 | 0.17 | 284 | 0 |
| classical monocytes | 0.13 | 0.2 | 284 | 0 |
| elicited macrophage | 0.08 | 0.25 | 284 | 0 |
| non-classical monocytes | 0.13 | 0.25 | 284 | 0 |
| pulmonary alveolar type 1 | 0.21 | 0.25 | 284 | 0 |
| pulmonary alveolar type 2 | 0.2 | 0.17 | 284 | 0 |
| capillary endothelial | 0.17 | 0.21 | 284 | 0 |
| ciliated columnar tracheobronchial tree | 0.28 | 0.14 | 284 | 0.33 |
| CD1c-pos | 0.1 | 0.3 | 284 | 0 |
| nasal mucosa goblet | 0.13 | 0.2 | 284 | 0 |
| vein endothelial cell | 0.23 | 0.17 | 284 | 0 |
| respiratory basal | 0.14 | 0.26 | 284 | 0 |
| pulmonary artery endothelial | 0.15 | 0.23 | 284 | 0 |
| alveolar type 1 fibroblasts | 0.25 | 0.19 | 284 | 0 |
| adventitial fibroblasts | 0.2 | 0.21 | 284 | 0 |

| Model | Significant perturbations | | | | SensX |
|---|---|---|---|---|---|
| | n_s | Delta | tau_a | tau_r | n_w |
| Synthetic | 2000 | lin(0.02, 1, 50) | 0.1 | 0.1 | multiple |
| ViT | 1000 | lin(0.02, 1, 50) | 0.1 | 0.1 | 500 |
| single-cell | 1000 | geom(0.0001, 1, 50) | 0.1 | 0.1 | 200 |
